# Potassium-chelating drug sodium polystyrene sulfonate enhances lysosomal function and suppresses proteotoxicity

**DOI:** 10.1101/2021.10.21.465344

**Authors:** Cyrene Arputhasamy, Mark Lucanic, Anand Rane, Minna Schmidt, Theo Garett, Anna C. Foulger, Michael Broussalian, Elena Battistoni, Rachel B. Brem, Gordon J. Lithgow, Manish Chamoli, Julie K. Andersen

## Abstract

Lysosomes are crucial for degradation and recycling of damaged proteins and cellular components. Therapeutic strategies enhancing lysosomal function are a promising approach for aging and age-related neurodegenerative diseases. Here, we show that an FDA approved drug sodium polystyrene sulfonate (SPS), used to reduce high blood potassium in humans, enhances lysosomal function both in *C*.*elegans* and in human neuronal cells. Enhanced lysosomal function following SPS treatment is accompanied by the suppression of proteotoxicity caused by expression of the neurotoxic peptides Aβ and TAU. Additionaly, treatment with SPS imparts health benefits as it significantly increases lifespan in *C*.*elegans*. Overall our work supports the potential use of SPS as a prospective geroprotective intervention.

**HIGHLIGHTS:**

- Sodium polystyrene sulfonate improves pH-dependent processing of lysosomal cargo, enhances proteotoxic stress resistance and extends lifespan in *C. elegans*
- Sodium polystyrene sulfonate boosts lysosomal function in human neuronal cells and reduces level of aggregation-associated phosphorylated-TAU

## INTRODUCTION

Lysosomal dysfunction is associated with aging and many age-related pathologies including Alzheimer’s disease (Bonam, Wang et al. 2019). The primary role of lysosomes is to carry out degradation and recycling of damaged proteins and other cellular components. The ability of lysosomes to act in response to multiple signaling inputs involving nutritional status, proteotoxic stress resistance, development and differentiation makes it a critical regulator of organismal survival (Lamming and Bar-Peled 2019, Savini, Zhao et al. 2019, Villegas, Lehalle et al. 2019). In fact, functional decline in lysosomal activity has been shown to severely affect lifespan and health span in many organisms. Conversely, interventions boosting lysosomal function are emerging as a potent means of promoting lifespan extension and delaying disease pathologies, particularly in relation to the central nervous system (Zhang, Sheng et al. 2009). Survival of neuronal cells that are destined to last throughout the life span relies heavily on lysosomal-based cellular recycling mechanisms. In this regard, identifying novel compounds and strategies to repurpose existing drugs to boost lysosomal function could help delay aging and age-related neurodegeneration.

Potassium restriction as a means to boosts vacuolar acidity and extend lifespan in *Sachromyces cerevisiae* was recently shown by Sasikumar et al (Sasikumar, Killilea et al. 2019). The lifespan extending effect of potassium restriction were also recapitulated by the supplementation with sodium polystyrene sulfonate (SPS), a potassium-chelating drug. The drug has been in medical use to treat hyperkalemia (high potassium levels) since 1958 (Hagan, Farrington et al. 2016). In the present study, we tested the functional conservation of SPS efficacy in higher model organisms. We specifically investigated effect of SPS supplementation on lysosomal function and proteotoxicity. Our results demonstrate that SPS enhances lysosomal function both in *C*.*elegans* and human neuronal cells and suppresses proteotoxicity caused by amyloid-β and hyper-phosphorylated-TAU, key drivers of Alzheimer’s disease. Overall our work supports the potential use of SPS as a prospective geroprotective intervention with the ability to suppress proteotoxicity caused by neurotoxic proteins and its future testing in preclinical mouse models.

## RESULTS

Earlier work demonstrated that SPS extends lifespan in yeast by boosting vacuolar acidity. We sought to test whether this mechanism of SPS is conserved and if it could also trigger pH changes in *C. elegans* lysosomes. To explore this idea, we first assayed lysosomal acidity with the dye acridine orange whose emission wavelength shifts when it is sequestered within acidic lysosomes (Moriyama, Takano et al. 1982). Results revealed a 4.5-fold increase in acridine orange staining in animals treated with SPS relative to untreated controls (Figure 1A). To evaluate the impact of SPS on lysosome function, we used a reporter strain expressing the LGG-1 lysosomal cargo protein fused to mCherry and an acid-inactivated GFP; quenching of GFP signal from this reporter reflects its delivery into the acidic lysosome (Chang, Kumsta et al. 2017). This strain, when treated with SPS, exhibited a 3-fold increase in lysosomal processing of the reporter relative to untreated controls (Figure 1B). For an independent test of lysosomal function, we made use of an assay quantifying sensitivity to the lysosomotropic agent chloroquine (Wibo and Poole 1974). After chronic SPS treatment, animals were 2.5-fold more resistant to a lethal chloroquine dose than untreated controls (Figure 1C). Taken together, these data point to a marked boost in lysosomal acidity and function in *C. elegans* in response to SPS.

**Figure 1.**
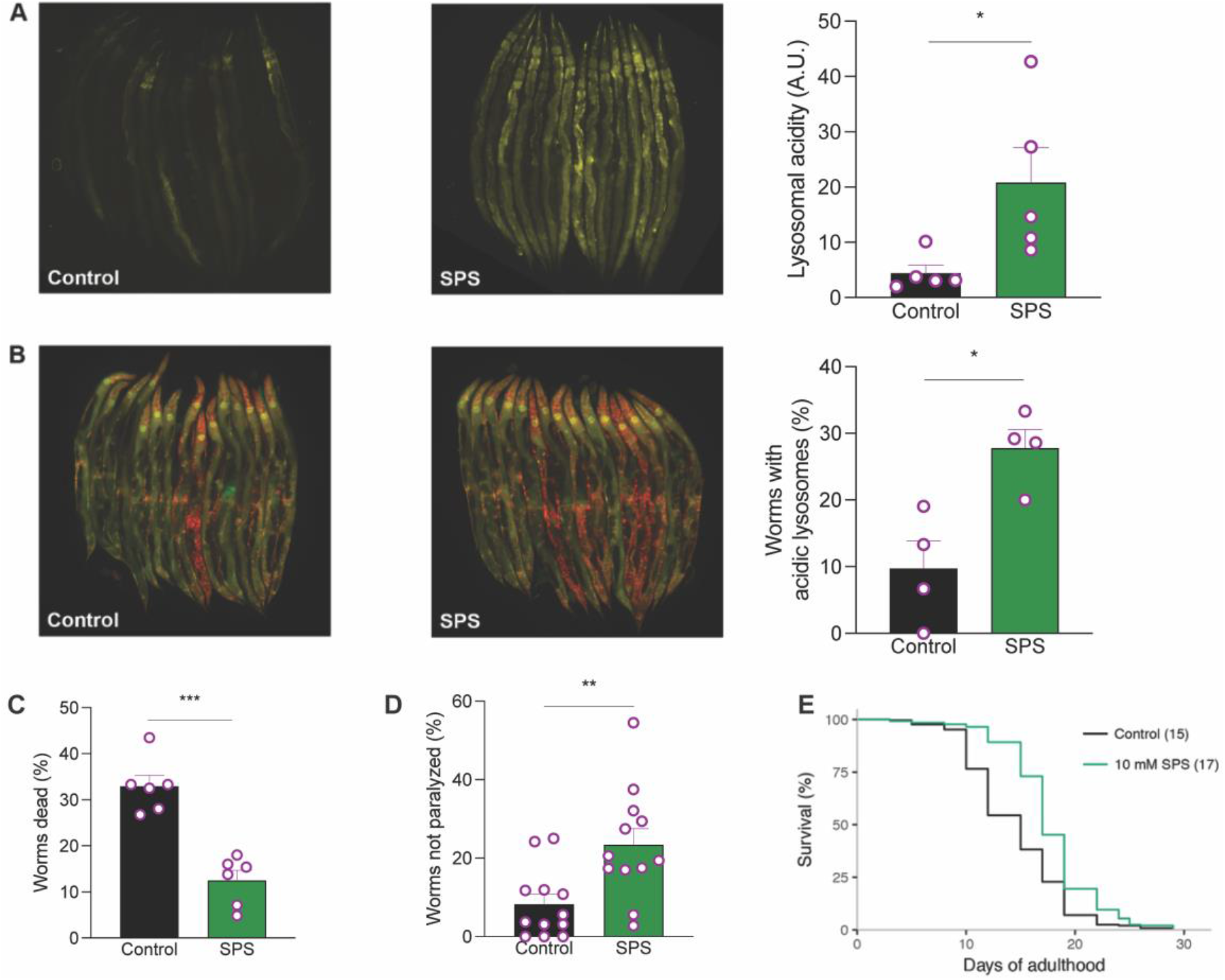
Sodium polystyrene sulfonate (SPS) treatment of worms improves pH-dependent processing of lysosomal cargo, enhances proteotoxic stress resistance and extends lifespan in *C. elegans*. **(A)**, Left, representative fluorescence microscopy images of wild-type worms maintained in the absence (control) or presence of 10 mM SPS and then treated with acridine orange to stain acidic lysosomes. Right, each bar reports mean acridine orange intensity per worm (*n* = 5 replicates, 15 worms per replicate) in arbitrary units (A.U.). **(B)**, Left, representative fluorescence microscopy images of worms maintained in the absence (control) or presence of 10 mM SPS expressing a fusion of the lysosomal cargo protein LGG-1 to mCherry and acid-inactivated GFP. Right, each bar reports the proportion of worms (*n* = 4 replicates, 15 worms per replicate) expressing the LGG-1 reporter that exhibited reduced GFP intensity relative to mCherry, reflecting processing of the reporter in the acidic lysosome. **(C)**, Each bar reports the mean viability (*n* = 6 replicates, 50 worms per replicate) of wild-type worms treated with the lysosomotropic agent chloroquine (20 mM), after maintenance in 10 mM SPS or control treatment. **(D)**, Each bar reports the mean proportion (*n* = 12 replicates, 35 worms per replicate) of amyloid-β- expressing worms maintained in the absence (control) or presence of 10 mM SPS that exhibited no detectable paralysis. Error bars report standard error. ***, one-tailed *t*-test *p* < 0.001; **, *p* < 0.01; *, *p* < 0.05; tests were paired for A and B and unpaired for C and D. **(E)**, Shown are results from a representative trial of a lifespan assay of wild-type *C. elegans* hermaphrodites on NGM plates (control) or NGM plates supplemented with 10 mM SPS. Numbers in parentheses report median lifespan in days (*n* = 262 for control, *n* = 268 for SPS). Significance of a Cox proportional hazards test comparing the two conditions was *p* = 3.5×10^−11^. Results from additional trials are reported in the supplementary table 1.

On the premise that SPS promotes the degradative function of lysosomes, we expected that this drug would also improve clearance of proteotoxic aggregates. To test this, we used a well-characterized *C. elegans* disease model in which expression of the aggregation-prone Aβ_1-42_ fragment of the human amyloid precursor protein in body wall muscle leads to temperature-dependent paralysis (McColl, Roberts et al. 2012). In this strain, SPS treatment conferred a 3-fold resistance to paralysis relative to controls (Figure 1D). We concluded that SPS helps resolve proteotoxic stress, mirroring the increased lysosomal function in these animals. Decline in lysosomal function and inability to maintain pH gradient is associated with accelerated aging in *C*.*elegans* (Anand, Holcom et al. 2020, Sun, Li et al. 2020). Thus, we tested ability of SPS to delay aging and extend lifespan in *C. elegans*. We observed WT animals treated with 10 mM SPS throughout adulthood lived 13% longer than controls (Figure 1E). We did not observe any general toxic effects of the SPS as there was no changes in the fecundity i.e. total number of eggs hatched (Fig. S1). Control experiments revealed no consistent effects of SPS on body movement (Figure S2), arguing against a feeding defect as implicated in the longevity of animals with compromised potassium transport (Davis, Somerville et al. 1995, Hamilton, Dong et al. 2005). Together, our data show that SPS has measurable health benefits in *C. elegans* in the context of physiology and behavior as well as increased viability into old age.

We next tested the ability of SPS to boost lysosomal function in mammalian neuronal cells. We exposed human H4 neuronal cells to SPS for 6 hours and quantified mRNA expression of lysosomal function-specific genes. Treatment of cells with 10 mM SPS significantly enhanced expression of several key lysosomal-specific genes up to levels of 1.5 - 2-fold, suggesting an increase in lysosomal function (Figure 2A). To further evaluate lysosomal function, we monitored the turnover rate of LC3-I, the mammalian homolog of *C. elegans* LGG-1, in SPS-treated neuronal cells both in the presence and absence of the lysosomal V-ATPase inhibitor bafilomycin A1. We observed that samples treated with SPS displayed increases in LC3-II levels which were further enhanced in the presence of bafilomycin A1 (Figure 2B and 2C). These results demonstrate that SPS treatment enhances lysosomal processing of LC3-II, confirming enhanced autophagy flux and lysosomal function. We further verified lysosomal function by performing a Dye Quenched-Bovine Serum Albumin (DQ-BSA) based assay (Marwaha and Sharma 2017). DQ-BSA is a heavily labeled fluorescent substrate of the lysosomal proteases, which shows reduced fluorescence under basal conditions due to a strong quenching effect. Upon hydrolysis of the DQ-BSA to single, dye-labeled peptides by lysosomal proteases, quenching is prevented, producing brightly fluorescent products. We observed that SPS-treated rat N27 neuronal cells showed a dose-dependent increase in fluorescence intensity (Figure 2D). This increase in fluorescence intensity was completely suppressed by the lysosomal inhibitor bafilomycin A1 (Figure 2D), demonstrating SPS enhances lysosomal function in mammalian neuronal cells. To test our hypothesis that SPS-enhanced lysosomal function suppresses proteotoxicity, we utilized human neuronal SH-SY5Y cells expressing full-length 4R-isoform of TAU. We found that SPS treatment reduced the level of aggregation associated phosphorylated-TAU (Figure 2E and 2F). Taken together, these results suggest a conserved function of the potassium chelator SPS in human neuronal cells and its therapeutic potential in suppressing proteotoxicity.

**Figure 2.**
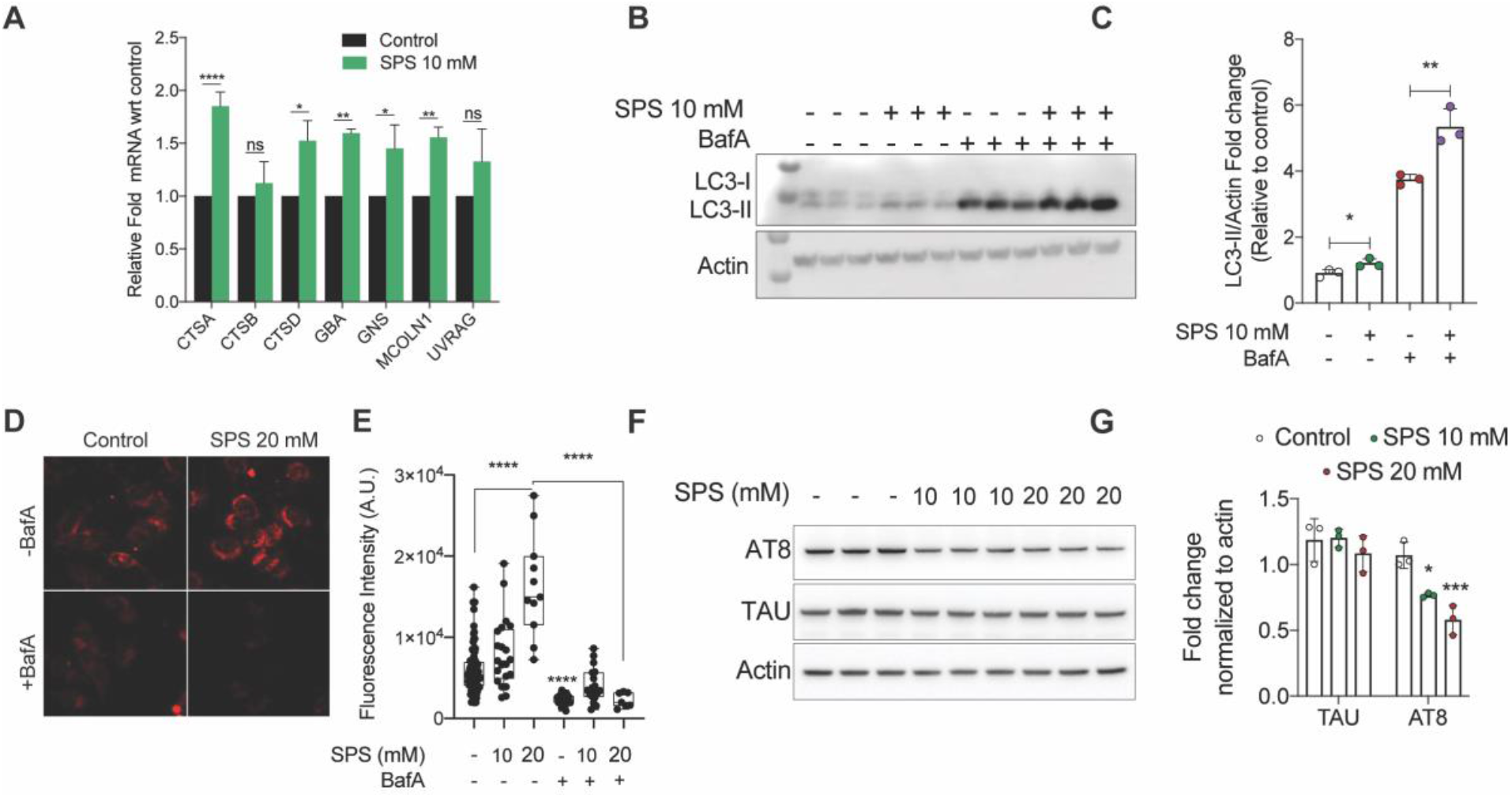
Sodium polystyrene sulfonate (SPS) boosts lysosomal function in mammalian neuronal cells and reduces level of aggregation-associated phosphorylated-TAU. **(A)**, Fold mRNA levels of lysosomal-specific genes in 10 mM SPS-treated human H4 neuronal cells relative to control samples (no SPS). Graph represents the mean ± SD of two biological repeats. *p*-values calculated using two-way ANOVA with Sidak’s multiple comparison test. **(B)**, Levels of LC3-I and LC3-II protein in 10 mM SPS-treated H4 neuronal cells in the absence and presence of the lysosomal V-ATPase inhibitor bafilomycin A1 (n=3 per condition). **(C)**, Fold changes in levels of normalized LC3-II protein relative to control sample (no SPS) quantified using NIH image J software. Graph represents mean fold intensity ± SD of Quantification of band intensity. *p*-values calculated using unpaired Student’s *t*-test. **(D)**, Represntative image of Dye Quenched-Bovine Serum Albumin (DQ-BSA) stained rat N27 neuronal cells. Images were taken in Axio Zeiss microscope using RFP filter. **(E)**, Graph represents mean fluorescence intensity ± SD of SPS treated (10 and 20 mM) rat N27 dopaminergic cells with and without addition of the lysosomal V-ATPase inhibitor bafilomycin A1. Each dot represents individual cell (n=3). *p*-values calculated using one-way ANOVA with Tukey’s multiple comparison test. **(F)**, Levels of TAU and phosphorylated TAU (Ser202, Thr205)/AT8 in 10 and 20 mM SPS-treated TAU expressing human SH-SY5Y neuronal cells (n=3 per condition). **(G)**, Fold changes in levels of normalized TAU and phosphorylated-TAU relative to control sample (no SPS) quantified using NIH image J software. Graph represents mean fluorescence intensity ± SD of SPS treated (10 and 20 mM) TAU expressing SH-SY5Y neuronal cells. *p*-values calculated using two-way ANOVA with Tukey’s multiple comparison test. *p*-values represents *p* < 0.001; **, *p* < 0.01; *, *p* < 0.05

## DISCUSSION

Our work demonstrates a previously unknown role for SPS in suppressing proteotoxicity caused by human neurotoxic proteins. The conserved role of SPS in enhancing lysosomal function in yeast, worms and human neuronal cells strongly suggests its likelihood of working in higher *in vivo* models. These findings are in line with other published work showing many potent anti-aging interventions are closely related to enhanced lysosomal function (Carmona-Gutierrez, Hughes et al. 2016). The lifespan-extending property of SPS further implies that the protective effect of SPS could potentially extend beyond suppression of proteotoxicity to delaying aging. The significance of our work is broad and is of general interest considering SPS is already used as a medication in humans and has been on the market for more than 60 years.

Although our study establishes that SPS enhances lysosomal function, it’s still not clear how exactly it does so. Due to its ability to disrupt ionic imbalance across the lysosomal membrane, SPS likely leads to lysosomal acidification. In mammalian cells the Na^+^/K^+^ ATPase pump regulates the sodium gradient across the membrane, which in turn drives the Na^+^/H^+^ antiporter and thus cellular pH (Kaplan 2002). SPS is a cross-linked polymer with a reactive sulfonic group that exchanges existing sodium (Na^+^) for potassium (K^+^) cations. This could potentially lead to a decrease in potassium concentration with simultaneous increases in the sodium concentration upon SPS exposure (Nepal, Bucaloiu et al. 2010). Overall, these changes by SPS profoundly affect lysosomal pH and function, imparting beneficial effects. Previous work in yeast cells shows that similar effects can be recapitulated by decreasing potassium concentration in the media, while no changes were observed by sodium supplementation, thus arguing in favor of potassium restriction as a potential mechanism resulting in a boost of lysosomal pH (Sasikumar, Killilea et al. 2019). Also, a recent work show that potassium starvation induces autophagy in yeast (Rangarajan, Kapoor et al. 2020), which falls in line with our observations, although these studies still needs to be validated in other organisms. Nevertheless, the idea of utilizing pharmacological compounds that alter Na^+^/K^+^ ionic imbalances to target neurodegenerative diseases is fairly novel and relatively untested. This is particularly interesting since several neurodegenerative diseases including Alzheimer’s have been reported to have a significant Na^+^/K^+^ ionic imbalance (Vitvitsky, Garg et al. 2012). Future experiments quantifying concentration of different ions in response to SPS treatment will be important to help decipher how SPS effect’s lysosomal function and suppresses proteotoxicity.

**Supplementary Figure 1:**
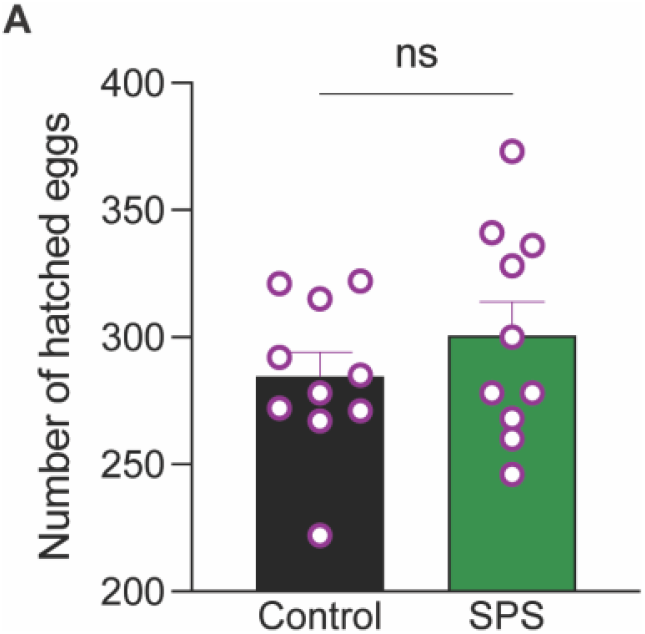
No changes in the fecundity upon SPS treatment. Total fecundity quantification i.e., total number of hatched eggs in N2 treated control or 10 mM SPS. *p-*value calculated using an unpaired Student’s t-test

**Supplementary Figure 2:**
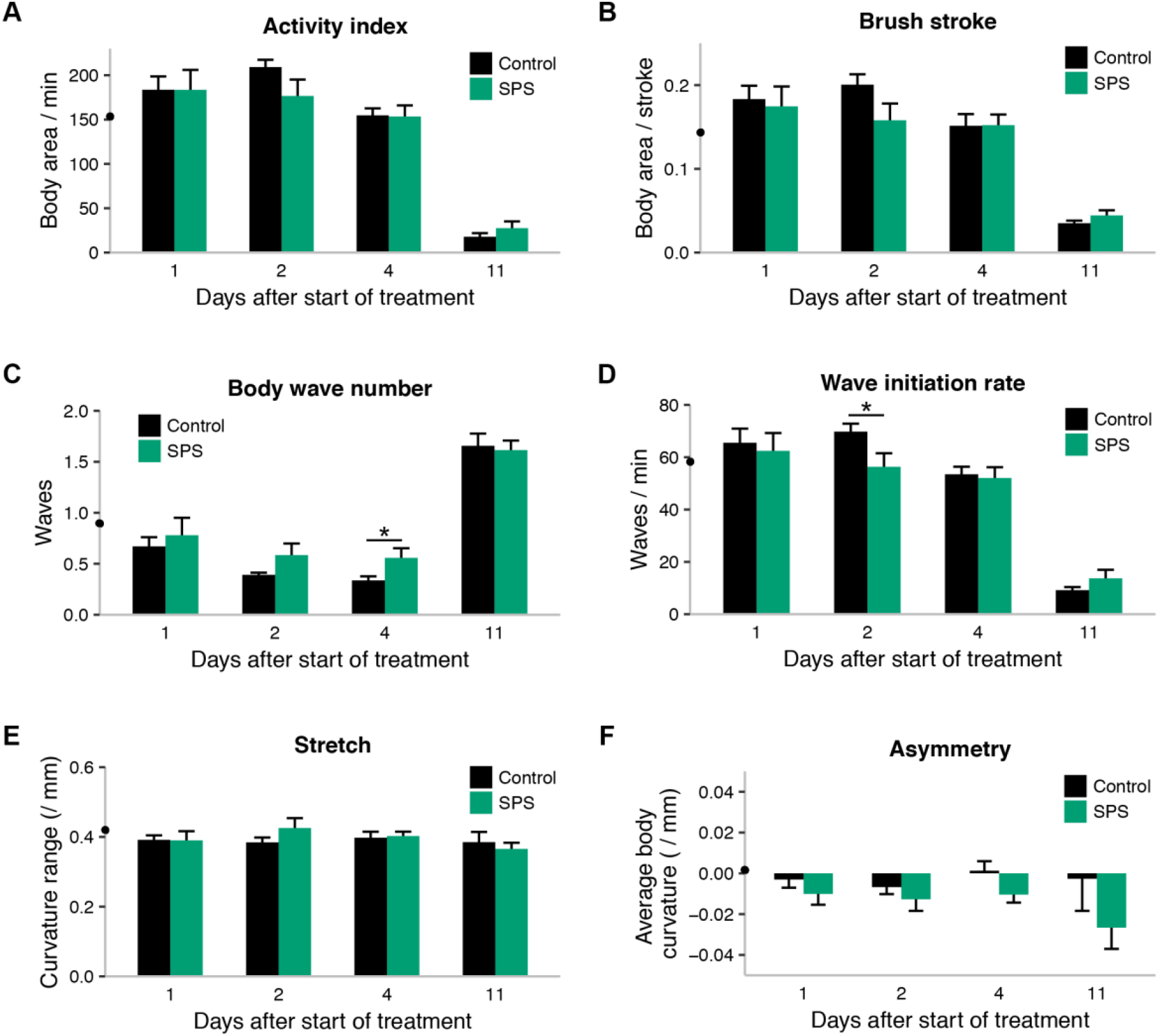
SPS has a negligible effect on worm movement. Each panel reports one attribute of the movement of wild-type worms maintained in the absence (control) or presence of 10 mM SPS over the indicated time course starting at day 1 of adulthood. **(A)**, The *y*-axis reports the number of pixels covered by a given animal over the time it took to undertake two swimming strokes. **(B)**, The *y*-axis reports the number of pixels covered by a given animal during a single swimming stroke. **(C)**, The *y*-axis quantifies the sinusoidal movement of the animals. Low scores (between 0-1) indicate that a single wave is propagating across the animal; higher scores indicate less coordinated movement. **(D)**, The *y*-axis reports the number of initiated body bends per minute. **(E)**, The *y*-axis reports the depth of bending of the swimming animal. **(F)**, The *y*-axis reports how balanced the swim posture was per stroke. In a given panel, each bar height reports the mean value of the respective measurement in worms of the respective treatment and timepoint, and error bars report standard error; the black point on the *y*-axis reports the mean value among 1-day-old adult animals pre-treatment. *, *t*-test 0.01 ≤ *p* ≤ 0.05.

**Supplementary Table 1:** Sodium polystyrene sulfonate (SPS) reproducibly extends worm lifespan. Each row reports results from one trial of lifespan assays of wild-type *C*.*elegans* on NGM plates (control) or NGM plates supplemented with 10 mM SPS. The last column reports results of a Cox proportional hazards test comparing the lifespans of treated and untreated animals in the respective trial.

| Trial | Treatment | No. of worms | Median lifespan (days) | *p* |
| --- | --- | --- | --- | --- |
| 1 | Control | 181 | 17 | 2.30E-02 |
|  | SPS | 57 | 19 |  |
| 2 | Control | 249 | 14 | 3.60E-11 |
|  | SPS | 248 | 16 |  |
| 3 | Control | 262 | 15 | 1.90E-11 |
|  | SPS | 268 | 17 |  |

## EXPERIMENTAL PROCEDURES

### Nematode culture, strains, and maintenance

Except where noted below, *C. elegans* hermaphrodites were maintained on nematode growth medium (NGM) agar plates seeded with *E. coli* bacterial strain OP50 at 20°C as described previously (Brenner 1974). Strains used in this study were wildtype Bristol N2; GMC101 dvIs100 [unc-54p::A-beta-1-42::unc-54 3’-UTR + mtl-2p::GFP]; and MAH215 sqIs11 [lgg-1p::mCherry::GFP::lgg-1 + rol-6] rollers.

### Worm sodium polystyrene sulfonate (SPS) treatment

A 500 mM stock of SPS was prepared in sterile water and stored at 4°C for up to 10 days. From the stock solution, 130 μl of the working SPS solution (10 mM) was prepared by mixing 60 μL of stock solution with 70 μL of sterile water, and was added to the top of the 35 mm NGM plates (3 mL NGM agar) already seeded with a bacterial OP50 lawn. SPS was distributed over the entire plate surface and allowed to dry in a sterile hood with the lid open for at least 1 hour. Plates were then allowed to sit at 20°C for 24 hours before use or before moving into 4°C for storage up to 2 weeks. For all worm experiments, animals were treated as follows except where noted below: a synchronous population was obtained by a 2-hour egg lay from gravid adult hermaphrodites, after which the adults were removed and the eggs were left to develop into adults at 20°C. On their first day of adulthood, the population was split into two, with half transferred to 35 mm NGM assay plates containing 10mM SPS and the other half transferred to control plates.

### Worm lifespan assay

Wild-type worms were synchronized and SPS-treated on day 1 of adulthood as above, except that following day 1, transfers continued periodically from each respective plate to a fresh plate of the same formulation (10 mM SPS or no-drug control) to ensure ample supply of drug and food. Survival was quantified by counting dead and live worms once every two or three days. We refer to the pipeline from egg lay through treatment and survival quantification as a trial, with a given such experiments were performed using drug-treated and control conditions in parallel and 35–40 worms on each of at least 3 replicate plates for each condition; three such independent trials were performed.

### Worm movement assay

Wild-type worms were synchronized and SPS-treated on day 1 of adulthood as above, except that following day 1, transfers continued periodically from each respective plate to a fresh plate of the same formulation (SPS or no-drug control) to ensure ample supply of drug and food. After 24h, 48h, 96h or 264h, worms were removed from their agar plate cultures and transferred into a swimming buffer. 30-second videos were then immediately captured of the swimming animals, which were then removed from the study and subjected to the CeleST analysis pipeline (Restif, Ibanez-Ventoso et al. 2014). ∼20 animals were captured for each time point and condition.

### Total fecundity quantification

The average number of fertilized eggs (fecundity) laid by worms was determined by transferring individual L4 larval staged wild-type worms onto NGM agar treated control or SPS bacterial plates (n = 9-10 worms per condition). Worms were allowed to lay eggs for the next 24 hours and then transferred to a fresh treated NGM agar bacterial plate. Worms were subsequently moved to fresh plates every day until they ceased laying eggs. On each plate, the total numbers of eggs laid and hatched were scored. The graph represents number of eggs hatched in each condition.

### Worm acridine orange (AO) staining

Wild-type worms were synchronized and SPS-treated on day 1 of adulthood as above, except that both SPS and control plates were supplemented with 10 μg/mL 5-fluoro-2′-deoxyuridine (FUdR) to inhibit progeny production. On day 2 of adulthood, worms were transferred from their respective plate to a fresh plate of the same formulation (SPS or no-drug control). On day 4, worms from these plates were transferred using a platinum pick to a 20 μl drop of AO (0.05 mg/ml in S-basal buffer, http://www.wormbook.org/chapters/www_strainmaintain/strainmaintain.html) on the side of a 2% agarose pad slide and allowed to stain for 15 minutes. After 15 minutes, the AO solution was removed using a pipette, and worms were washed with S-basal buffer. ∼15 worms were then transferred to a drop of fresh S-basal buffer on the agarose pad, to which was added 5 μl of 2 mM levamisole to immobilize. Worms were visualized under a rhodamine filter on a Zeiss fluorescence microscope. Intensity in a region delimited manually from the perimeter of each worm was quantified in ImageJ. The pipeline from growth through treatment, staining and quantification we refer to as a replicate, with a given such experiments were performed using drug-treated and control conditions in parallel; five such independent replicates were performed.

### Tandem-tagged LGG-1 reporter imaging

MAH215 worms were synchronized and SPS-treated on day 1 of adulthood as above. After two hours on SPS or no-drug control plates, ∼15 worms were imaged using GFP and rhodamine filters on a Zeiss fluorescence microscope. GFP and mCherry intensity from individual worms were quantified using ImageJ. For each worm, we tabulated the ratio of mCherry and GFP intensities and, as a metric of LGG-1 processing, we tabulated the proportion of worms where this ratio exceeded 1.5. The pipeline from growth through treatment, staining, and quantification we refer to as a replicate, with a given that such experiments were performed using drug-treated and control conditions in parallel; four such independent replicates were performed.

### Chloroquine toxicity assay

Wild-type worms were synchronized and SPS-treated on day 1 of adulthood as above, except that following day 1, transfers continued periodically from each respective plate to a fresh plate of the same formulation (SPS or no-drug control) to ensure ample supply of drug and food. On day 7, worms from each plate were transferred to 20 mM chloroquine supplemented NGM agar plates without SPS. On day 9 and day 11, worms were transferred to fresh chloroquine plates. On day 11, viability was quantified by counting dead and live worms. We refer to the pipeline from egg lay through SPS treatment, chloroquine treatment, and survival quantification as a trial, with a given such experiments were performed using SPS-treated and control conditions in parallel using ∼50 worms on each of 3 replicate plates for each condition. Two such independent trials were performed.

### Worm amyloid-beta paralysis assay

A synchronous population of GMC101 worms was obtained by 2-hour egg-lay at 20°C. At the L4 stage, 48-hours after egg-lay, the population was split into two, with half transferred to plates containing 10 mM SPS and the other half to control plates; all were incubated at 25°C for an additional 48 hours. At this timepoint, body movement was assessed as follows. A given worm was scored as paralyzed if it (1) failed to complete a full body movement, *i*.*e* a point of inflection traversing the entire body length, either spontaneously or touch-provoked using platinum wire, and (2) exhibited a halo of cleared bacteria around its head, indicative of insufficient body movement to access food. We refer to the pipeline from egg lay through SPS treatment and paralysis scoring as a trial, with a given that such experiments were performed using SPS-treated and control conditions in parallel and ∼35 worms on each of 4 replicate plates for each condition. Three such independent trials were performed.

### Cell culture and maintenance

Human H4 neuroglioma cells (ATCC) were maintained in Dulbecco’s Modified Eagle Medium (DMEM), human SH-SY5Y-GFP-Tau neuroblastoma cells (ATCC) were maintained in HyClone Classical Liquid DMEM-F12 media, and rat N27 dopaminergic neural cells (Millipore) were maintained in Roswell Park Memorial Institute (RPMI) 1640 medium. All culture media were supplemented with 10% FBS (Serum Plus – II, Corning) and 1% penicillin-streptomycin (Corning). Cells were incubated at 37ºC in a 5% CO_2_ humidified cell culture incubator.

### Chemicals and buffers

Sodium polystyrene sulfonate (SPS) (Sigma) was dissolved in the appropriate basal media and stored at 4°C for up to 10 days at a concentration of 1M. Dulbecco’s phosphate-buffered saline (DPBS) was purchased from Corning. PBST buffer (PBS plus 0.1% Tween-20) was used for blot washing. Blocking buffer and primary antibody incubation solution was 5% non-fat dairy milk powder (VWR) or 5% (w/v) BSA (Sigma) for phospho-antibodies.

### Cellular SPS treatment

The working SPS solution (10 mM) was prepared by mixing stock solution in complete media at a dilution of 1:100 and adding to adherent cells already seeded at the proper density in tissue culture plates. Cells were treated with SPS for 4-6 hours, with some samples being pretreated with bafilomycin-A1 (Sigma-Aldrich) for 1 hour at 100 nM.

### RNA isolation and qPCR

H4 cells were treated with SPS as described and collected for RNA isolation. RNA was isolated using a Zymo Research Quick-RNA miniprep kit (Zymo). Quality and concentration of RNA was confirmed on a NanoDrop 2000 spectrophotometer (Thermo). From the RNA isolation, 1ug was used to generate cDNA using a High-Capacity cDNA Reverse Transcriptase kit (Applied Biosystems). The qPCR was performed using 20 ng of cDNA product per reaction in the presence of Lightcycler 480 SYBR green dye (Roche). Lysosomal gene primers were designed using PrimerBLAST and ordered from Eurofins; they were used at a subsequent working concentration of 200 nM. The sequence of forward(F) and reverse (R) of the primers (5’-3’) used are: CTSA-F:CAGGCTTTGGTCTTCTCTCCA, CTSA-R: TCACGCA TTCCAGGTCTTTG, CTSB-F: AGTGGAGAATGGCACACCCTA, CTSB-R: AAGAAGCCATTGTCACCCCA, CTSD-F: AACTGCTGGACATCGCTTGCT, CTSD-R: CATTCTTCACGTAGGTGCTGGA, GBA-F: TGGGTACCCGGATGATGTTA, GBA-R: AGATGCTGCTGCTCTCAACA, GNS-F: CCCATTTTGAGAGGTGCCAGT, GNS-R: TGACGTTACGGCCTTCTCCTT, MCOLN1-F: TTGCTCTCTGCCAGCGGTACTA, MCOLN1-R: GCAGTCAGTAACCACCATCGGA, UVRAG-F: CATCTGTGTCTTGTTTCGTGG, UBRAG-R: TTCA TTTTGGTTTCGGGCATG

### DQ Red assay

N27 cells were seeded in a 4-well chamber slide at 30,000 cells per well. The next day, DQ Red BSA dye (ThermoFisher Scientific) was prepared at a working of concentration 10 μg/ml (1:200 dilution) in N27 media. Cells were washed once in pre-warmed DPBS, then DQ dye was added to cells followed by incubation for 1 hour. The DQ media was then removed and replaced with media +/- bafilomycin. After 1 hour, SPS was added at a final concentration of 10 mM for 4 hours. Cells were then fixed with 4% PFA prepared in DPBS for 10 minutes. Slides were mounted with Prolong Gold Antifade Mountant in the presence of DAPI (ThermoFisher Scientific). Images were taken on a confocal microscope (Zeiss) at 40x magnification.

### Western blotting

Cells treated with SPS either alone or in conjunction with bafilomycin-A1 were collected and subsequently lysed in lysis buffer (50 mM Tris-HCl pH 8, 150 mM NaCl, 1% NP-40) supplemented with PhosStop phosphatase inhibitor cocktail (Sigma Aldrich) and protease inhibitor cocktail (Sigma Aldrich). Protein concentrations were quantified by Bradford assay per manufacturer’s instructions (BioRad). Samples were normalized per protein via dilution in lysis buffer and SDS protein dye. Samples were heated at 90ºC for 15 minutes and then centrifuged at 15,000 xg for 10 minutes. Samples were next electrophoresed on a NuPAGE 4-12% Bis-Tris gel (ThermoScientific), the resulting gel transferred to PVDF membrane, blocked in blocking buffer (5% milk in PBST) and probed with primary and then secondary antibodies. The primary antibodies used were p-Tau (S422) from ThermoFisher (44-764G), p-Tau (S204, T205) from ThermoFisher (MN1020), Tau from ThermoFisher (MN1000), LC3B from Cell Signaling Technologies (2775S), and β-actin from Cell Signaling Technologies (3700S).

## ACKNOWLEDGEMENTS

The authors thank Dr. Arjun N. Sasikumar for helpful discussion. This work was supported by NIH RF1 AG057358 to JKA. The *C. elegans* strains used in this work were provided by the *Caenorhabditis* Genetics Center (CGC), funded by the NIH Office of Research Infra-structure Programs (P40OD010440). MC is supported by the postdoctoral fellowship from the Larry L. Hillblom Foundation.

## AUTHOR CONTRIBUTIONS

Conception and design, R.B.B., M.C. and J.K.A.; Data acquisition and analysis, C.A., M.L., A.R., M.S., T.G., A.F., M.B., E.B. and M.C.; Writing –Review & Editing, R.B.B., M.C., G.J.L. and J.K.A.; Supervision, M.C. and J.K.A.; Funding Acquisition, G.J.L. and J.K.A

## DATA AVAILABILITY STATEMENT

The data that support the findings of this study are available from the corresponding author upon reasonable request.

